# Multimodal evidence on shape and surface information in individual face processing

**DOI:** 10.1101/299933

**Authors:** Dan Nemrodov, Marlene Behrmann, Matthias Niemeier, Natalia Drobotenko, Adrian Nestor

## Abstract

The significance of shape and surface information for face perception is well established, yet their relative contribution to recognition and their neural underpinnings await clarification. Here, we employ image reconstruction to retrieve, assess and visualize such information using behavioral, electroencephalography and functional magnetic resonance imaging data.

Our results indicate that both shape and surface information can be successfully recovered from each modality but that the latter is better recovered than the former, consistent with its key role for face representations. Further, shape and surface information exhibit similar spatiotemporal profiles, rely on the extraction of specific visual features, such as eye shape or skin tone, and reveal a systematic representational structure, albeit with more cross-modal consistency for shape than surface.

Thus, the present results help elucidate the representational basis of individual face recognition while, methodologically, they showcase the utility of image reconstruction and clarify its reliance on diagnostic visual information.

## Introduction

The segregation of shape and surface information defines a fundamental aspect of visual processing and cortical organization (Livingstone & Hubel, 1988; Van Essen & Deyoe, 1995) both in the human (Cant, Large, McCall, & Goodale, 2008; Lafer-Sousa, Conway, & Kanwisher, 2016; Vinberg & Grill-Spector, 2008) and the monkey brain (Conway, Moeller, & Tsao, 2007). Accordingly, this distinction has played a prominent role in accounts of face recognition (Bruce & Young, 1998). Extensive research has documented the importance of both types of information in face perception (Biederman & Kalocsai, 1997, Jiang, Blanz, & O’Toole, 2006; O’Toole, Vetter, & Blanz, 1999; Russell et al., 2007; Russell & Sinha, 2007; Vuong, Peissig, Harrison, & Tarr, 2005), but the relative weight of shape and surface properties has been heavily debated, with either the former (Jiang, Blanz, & Rossion, 2011; Lai, Oruc & Barton, 2013) or the latter (Bruce et al., 1991; Bruce & Langton, 1994; Hole, George, Eaves, & Rasek, 2002; Kaufmann & Schweinberger, 2008; Russell, Sinha, Biederman & Nederhouser, 2006) considered dominant. Arguably, this debate arises from a lack of specificity in identifying the shape and surface features critical for individual face processing (Burton et al., 2015). Thus, the current research aims to uncover the nature of the information involved in individual face processing along with its accompanying neural profile.

To address the challenge above, here, we appeal to neural-based image reconstruction (Shen, Dwivedi, Majima, Horikawa & Kamitani, 2018; Miyawaki et al., 2008; Naselaris et al., 2009, Nishimoto et al., 2011; Thirion et al., 2006), namely, the endeavor of reconstructing the appearance of visual objects from neural activity prompted by their processing. While this endeavor has relied primarily on functional magnetic resonance imaging (fMRI), more recently, additional modalities have been used successfully as well. For instance, facial image reconstruction has been carried out using single-cell recordings (Chang & Tsao, 2017), electroencephalography (EEG) data (Nemrodov et al., 2018) and behavioral data (Chang et al., 2017; Zhan et al., 2017), in addition to fMRI (Cowen, Chun, & Kuhl, 2014; Lee & Kuhl, 2016; Nestor, Plaut, & Behrmann, 2016). Thus, in theory, image reconstruction can provide a powerful platform for investigating shape/surface processing in face individuation via multiple behavioral and neuroimaging modalities. Concretely, image reconstruction can be used to uncover, assess and compare facial shape and surface information recovered from distinct modalities.

To this end, we rely on data assessing individual face processing gleaned from behavioral (Nestor, Plaut & Behrmann, 2013), EEG (Nemrodov et al, 2018) and fMRI data (Nestor, Plaut & Behrmann, 2016). Specifically, for each modality, we aim to recover the shape and surface content of a common set of face stimuli as perceived by human observers (see Figure 1). In addition, the same procedure is conducted with an image-based theoretical observer (TO) allowing us to compare the informational content of multiple empirical and TO reconstructions.

**Figure 1.**
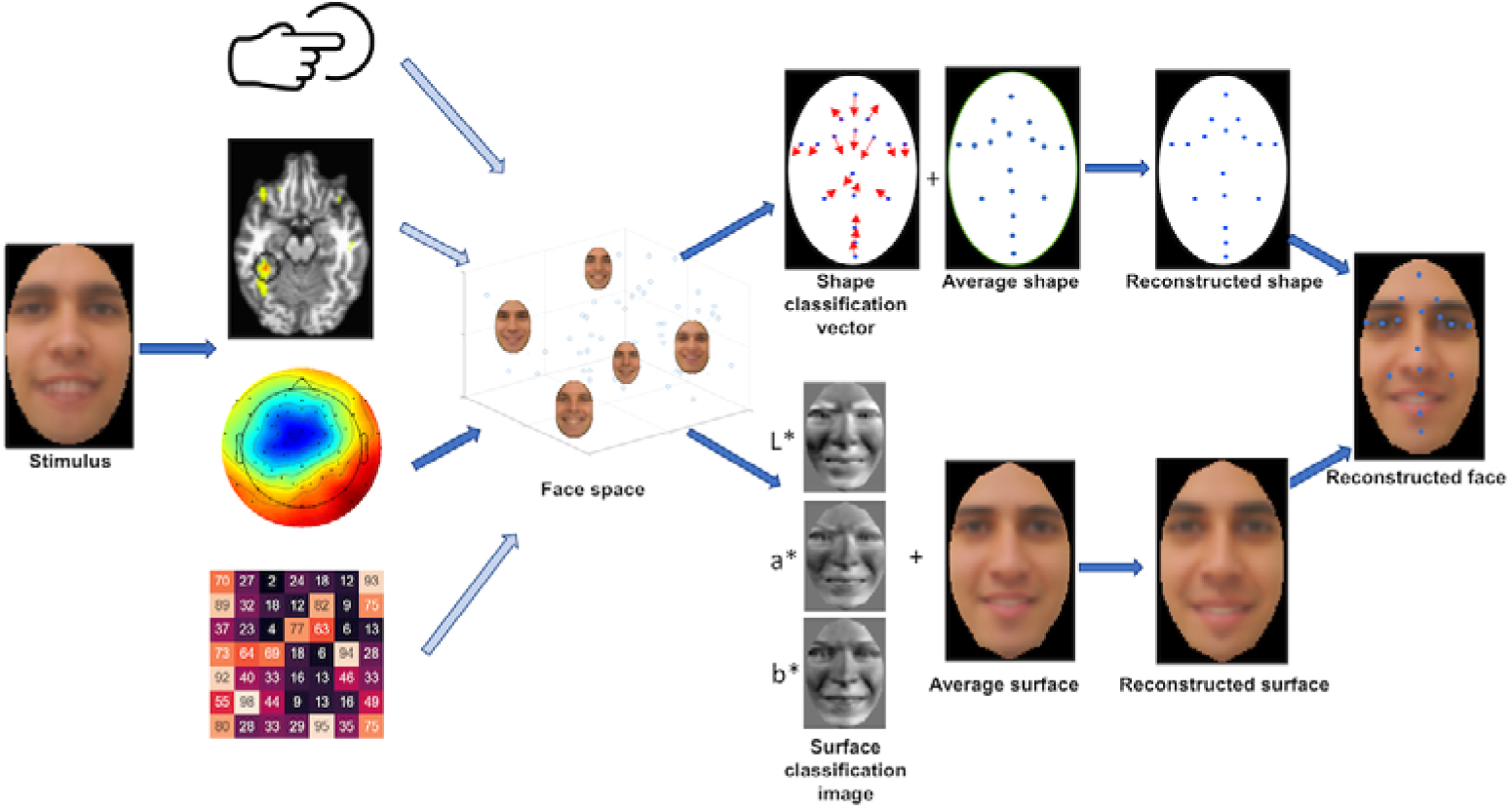
Schematic illustration of the reconstruction procedure. Behavioral, fMRI, EEG and TO data associated with viewing face stimuli support, separately, the estimation of a multidimensional face space (for convenience, a single example based on EEG data, indicated by the dark blue arrow, is shown, but similar results can be achieved from other modalities). Shape information and surface information are derived from the structure of this space and combined into facial image reconstructions (only a representative subset of fiducial points are displayed; L*, a* and b* correspond to the lightness, red-green and yellow-blue channels of color vision as encoded in CIEL*a*b*). Due to copyright restrictions all identifiable stimulus images were replaced with computer-generated images not used during experimental testing.

Accordingly, we appeal here to an influential approach for analyzing face images into shape and surface properties (Craw & Cameron, 1991; Kramer, Jenkins & Burton, 2016; Tiddemann, Burt & Perrett, 2001; Vetter & Troje, 1995). Specifically, this approach involves marking the positions of a set of fiducial points (e.g., the corners of the eyes or the tip of the nose) that deliver shape information. Then, faces are warped to a standard shape (i.e., a preset configuration of fiducial points) yielding ‘shape-free’ images that deliver surface information. To be clear, shape derived in this manner encompasses two sources of information: configural information, conceived as metric distances between different face parts (Maurer et al., 2002; Tanaka & Gordon, 2011), and local information associated with the geometric structure of specific face parts such as eye shape or mouth shape (Cabeza & Kato, 2000; Gold, Mundy & Tjan, 2012; Rakover, 2002). In contrast, surface contains information about the reflectance properties of a face (e.g., hue, specularity, albedo) that also play a role in individual face recognition (Hancock, Burton & Bruce, 1996; Russell et al., 2007; Taschereau-Dumouchel et al., 2010) – such information is alternatively referred to as ‘texture’, ‘pigmentation’ or ‘surface reflectance’.

The appeal to shape-surface decomposition allows us to address a number of related questions. First, can image reconstruction separately recover facial shape and surface information from different modalities and, if so, how well? Second, what is the spatiotemporal profile of shape and surface processing? Third, what specific shape/surface features are recovered through reconstruction? And forth, do different modalities reveal similar or complementary information about face representations? More generally, the present work evaluates and confirms the ability of a novel methodological paradigm to exploit multimodal evidence in the effort to elucidate the representational content of individual face processing.

In summary, the current work embarks on a comprehensive investigation of facial shape and surface processing by appeal to powerful and novel image-reconstruction methodology as applied to multimodal data. Accordingly, this work serves a twofold purpose by shedding light on the psychological and neural profile of facial shape/surface processing and by clarifying the informational content responsible for the success of image reconstruction.

## Results

Our investigation relies on the representational structure underlying facial processing as revealed by multiple modalities for a common set of stimuli. This structure allows us to relate the outcome of different modalities to each other as well as to derive shape and surface estimates of face representations. Such estimates can be assessed in terms of their reconstruction success and of their spatiotemporal neural profile. Further, such estimates can be recombined into image reconstructions approximating the visual appearance of the percepts associated with viewing specific face stimuli.

### Representational similarity

Estimates of pairwise face similarity were computed across 108 images (54 identities x 2 expressions, neutral and happy) for each of four data types: (i) behavioral, based on similarity ratings; (ii) EEG, based on neural discriminability across occipitotemporal (OT) electrodes; (iii) fMRI, based on neural discriminability across multiple fusiform gyrus (FG) areas, and (iv) TO, based on pixelwise image similarity. In particular, we note that fMRI estimates relied jointly on patterns of activation from four distinct FG areas (see Materials and Methods, Representational similarity analysis) - these areas were selected due to their ability to support face decoding in previous work (Nestor et al, 2016). For clarity, stimulus-specific multivoxel patterns were concatenated here across the four areas and subjected to pattern classification.

Next, Spearman rank correlations were computed for each pair of modalities across corresponding similarity estimates (i.e., 1431 pairwise estimates for 54 identities, averaged across expressions). Overall, we found that all data types correlated with each other (p<0.01, Bonferroni-corrected) providing initial evidence for common representational structure across modalities (Figure 2a).

**Figure 2.**
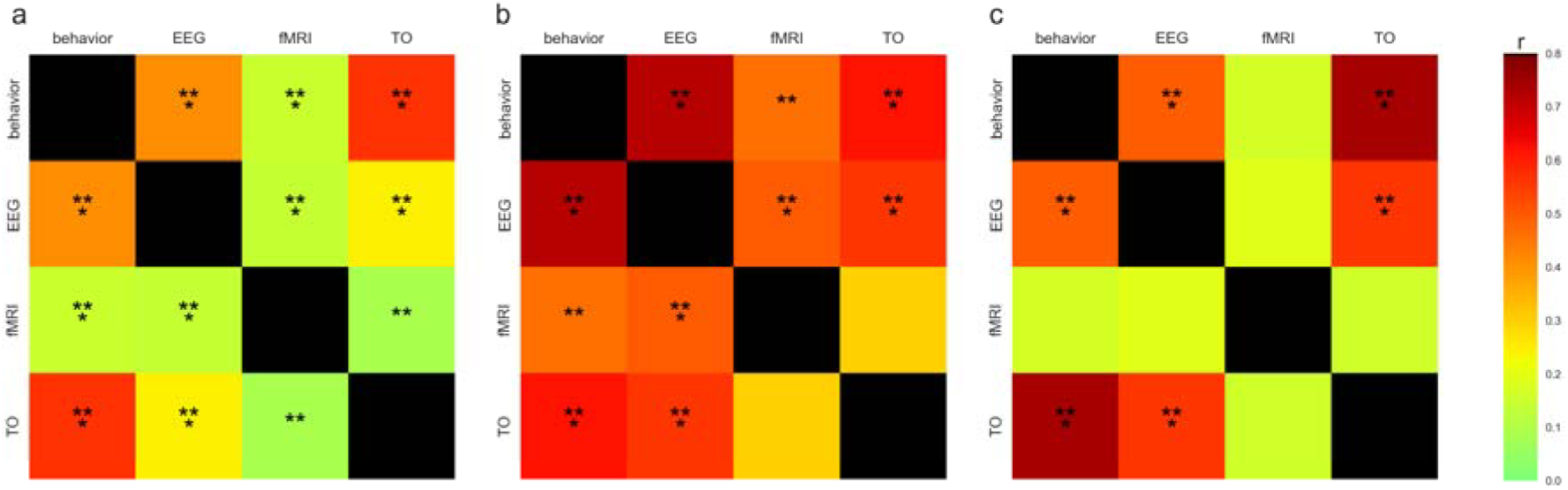
Correlations between different data types were based on: (a) pairwise face similarity/discriminability estimates (Spearman correlation across 1431 facial identity pairs); (b) shape and (c) surface reconstruction accuracy (Pearson correlation across 54 facial identities). All modalities are correlated with each other in terms of face similarity but only some in terms of shape and surface information (** p<0.01, *** p<0.001; Bonferroni correction across comparisons).

### Shape and surface reconstruction

Accuracy estimates of shape and surface reconstruction were separately computed for each data type (Figure 3). Critically, we found that all estimates were above chance for both facial expressions (permutation test, p<0.05, Bonferroni correction for 24 comparisons). Overall, for empirical modalities, surface reconstructions were more accurate than shape reconstructions (paired-comparison permutation test, p<0.01) with the exception of fMRI results for happy faces where the difference was only marginally significant (p=0.085). In contrast, TO results showed no difference in accuracy between shape and surface (p>0.151), suggesting that the reconstruction method can, in theory, retrieve the two types of information with equal success from the current stimulus sets. Further, no difference was found between neutral and happy faces for either shape or surface for any data type (p>0.104 for all other than fMRI-based shapes, in which case happy faces yielded marginally more accurate reconstructions than neutral ones, p=0.079).

**Figure 3 with 1 supplement.**
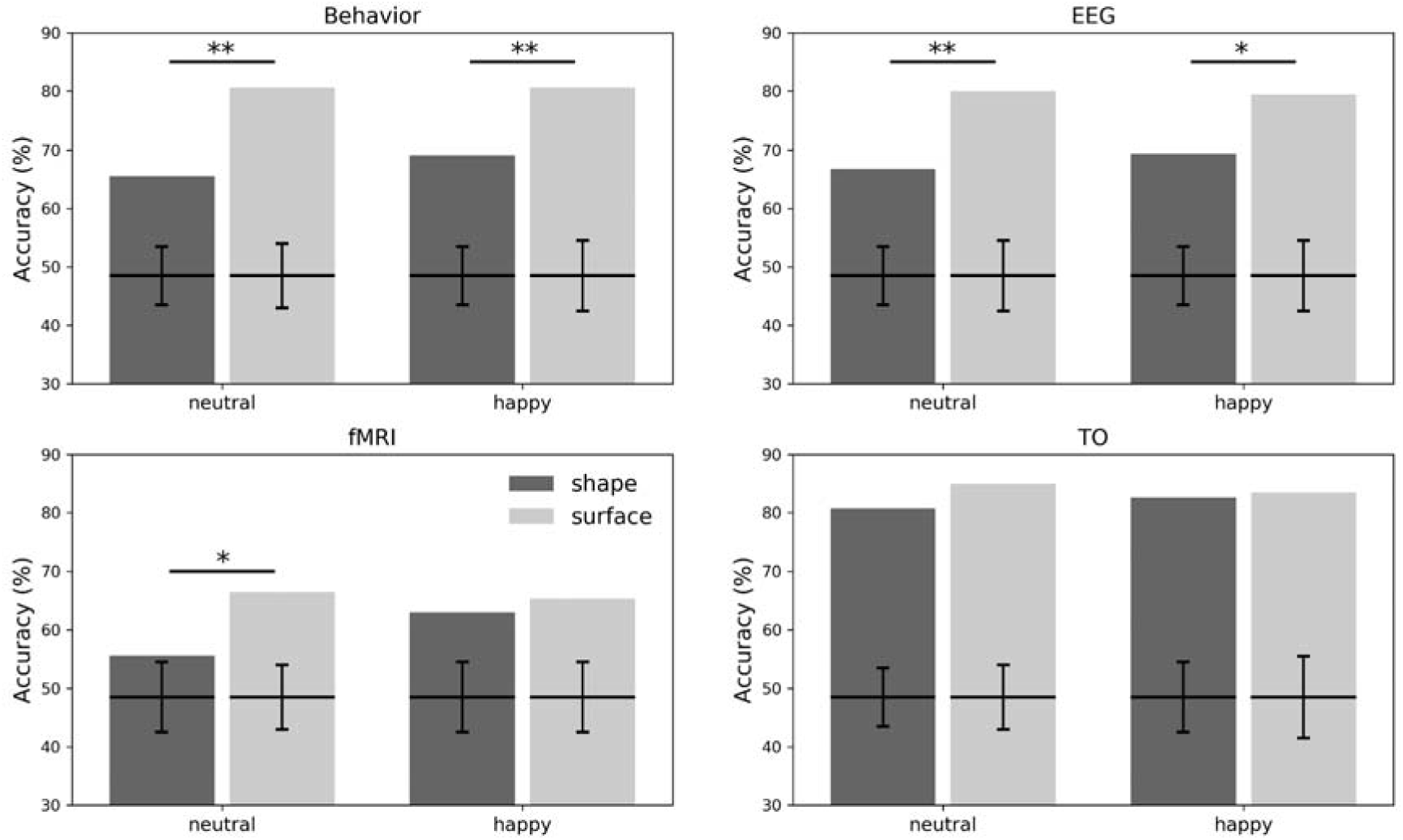
Shape and surface reconstruction accuracies for four modalities. All accuracy estimates are above chance (all *p*’s<0.01, Bonferroni-corrected; two-tailed permutation tests; confidence intervals based on 10^4^ shuffles of identity labels in the corresponding face space). Surface information was retrieved more accurately than shape information for empirical modalities (two-tailed paired-comparison permutation tests; * *p*<0.05; ** *p*<0.01) but not for TO. Figure 3 – Figure supplement 1. Accuracy estimates of surface reconstruction for each color channel. Results are shown for each modality collapsed across emotional expression. All estimates are above chance (all *p*’s<0.01, Bonferroni-corrected; two-tailed permutation tests; confidence intervals based on 10^4^ shuffles of identity labels in the corresponding face space). No difference was noticed across color channels for any modality (all *p*’s>0.05; two-tailed paired-comparison permutation tests).

Given the systematic advantage of surface over shape reconstructions, we proceeded to assess reconstruction accuracy separately for each color channel. This assessment is particularly relevant since informative shape-from-shading cues may be present as lightness patterns in what we refer to as ‘facial surfaces’ (Attick, Griffin & Redlich, 1996). Hence, the surface advantage noticed above could be due to another source of shape information rather than to genuinely ‘shape-free’ surface information. To evaluate this possibility, we estimated reconstruction accuracy for each color channel: we averaged such estimates across expression, given the absence of an expression effect above, and we compared them with each other separately for each modality. This analysis revealed that color components and lightness support equivalent levels of reconstruction accuracy for every modality (Figure 1 – figure supplement 1). More precisely, no difference was noticed between any two components (two-tailed paired-comparison permutation tests; p>0.05, uncorrected) ruling out the shape-from-shading hypothesis above.

Next, regarding the relative performance of different modalities, we note that fMRI seemed to perform more poorly than other data types. However, a direct comparison of different modalities in terms of accuracy may be misleading in that the corresponding experiments followed different protocols suitable for the corresponding modalities (e.g., different experimental tasks, different numbers of trials, different numbers of participants). At the same time, we note though that representational similarity analysis confirmed the presence of corresponding structure across data types. To further explore this correspondence in terms of shape and surface information, reconstruction accuracy, averaged across expressions, was correlated for each pair of data types (Figure 2b, c). This analysis found, in the case of shape, that empirical modalities all correlated with each other with the exception of fMRI and TO. However, in the case of surface properties, behavioral reconstructions were correlated with EEG and TO, but not with fMRI, pointing to potentially different surface information available in fMRI data.

To clarify the results above in the context of the relationship between brain and behavior, multiple linear regression was employed to account for behavioral accuracy based on all other data types. Separate analyses for shape and surface information both yielded significant models (shape: R^2^_adj_=0.55, *p*<0.001; surface: R^2^_adj_=0.47, *p*<0.001). In more detail, for shape information, EEG (*β*=0.41, p<0.001), fMRI (*β*=0.14, *p*=0.01) and TO (*β*=0.39, *p*<0.001) all provided significant independent contributions in accounting for behavior. In contrast, for surface information, only TO (*β*=0.64, *p*<0.001) and, marginally, EEG (*β*=0.14, *p*=0.075), were significant predictors of behavioral accuracy. Thus, different modalities appear to contain only partly overlapping information and to make distinct contributions in accounting for behavioral performance.

Next, to pinpoint the source of recovered information, accuracy was locally computed for each fiducial point, in the case of shape, and for each pixel and color channel, in the case of surface properties. Specifically, the coordinates of each fiducial point within a reconstructed image were compared relative to the corresponding point in the stimulus images and point-specific accuracy was estimated as the percentage of instances for which the Euclidean distance to the corresponding point in the target stimulus was smaller than to that in any other stimulus. Accuracy heatmaps averaged across all reconstructed shapes are displayed in Figure 4. This analysis revealed that the shape of the eyes was better recovered than other information for all empirical modalities. The same appeared to be the case for TO reconstructions; however, additional information regarding the shape of the mouth and the eyes was also recovered relatively well here.

**Figure 4.**
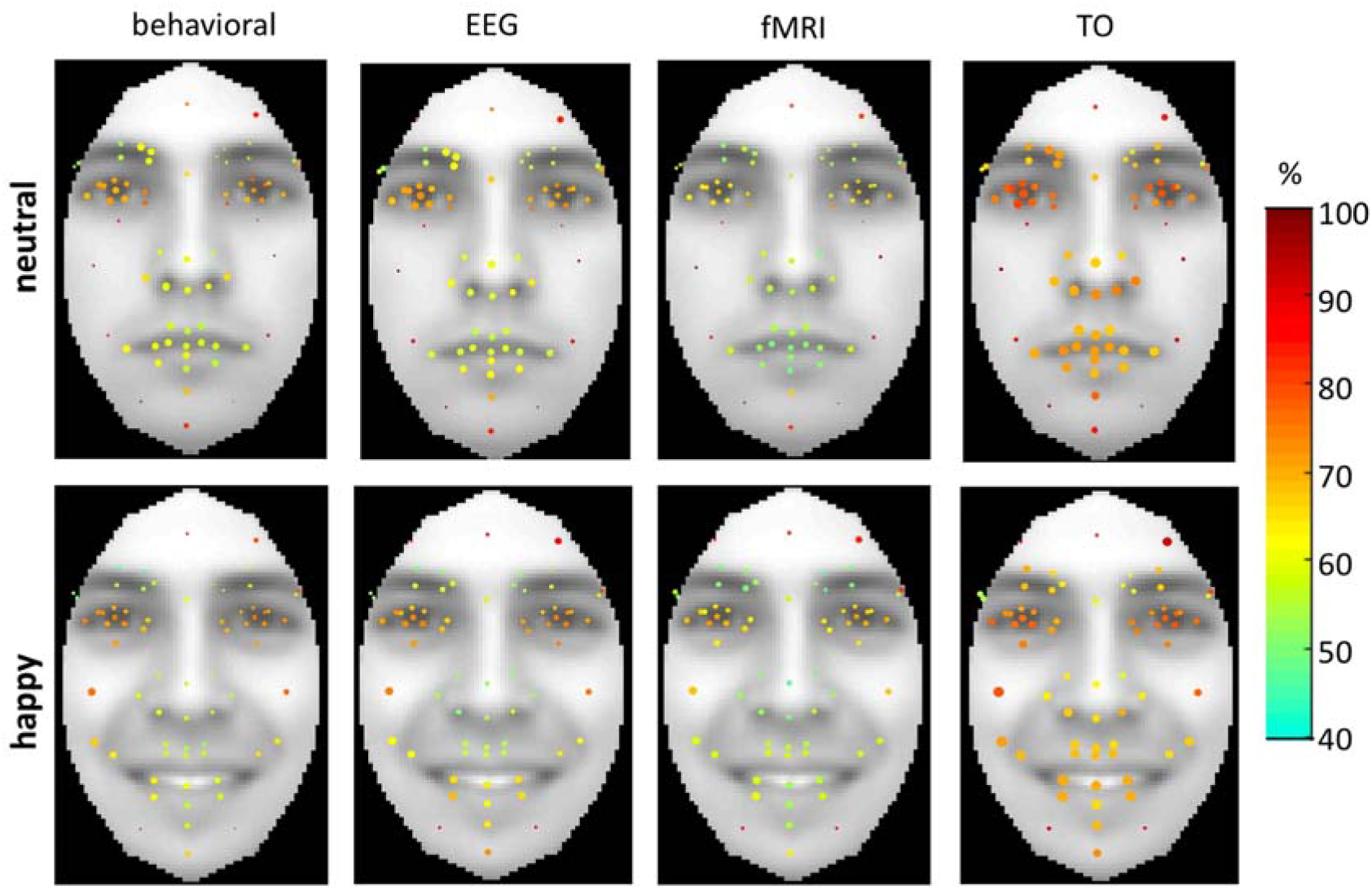
Accuracy heatmaps for shape reconstruction across fiducial points overlaid on average neutral and happy faces. The size of the circles is proportional with position variance across the face (i.e., larger circles indicate more variability in fiducial point position across different individual faces) while color indicates average reconstruction accuracy across 54 facial identities. Shape information is best approximated across the eyes though differences in both global and local accuracy can be noticed across modalities.

A similar procedure was followed for deriving surface heatmaps except that pixel intensity values (e.g., lightness as coded in the L* channel), rather than geometrical coordinates, were considered in this case (see Figure 5 and Figure 5 – supplement figure 1 for neutral and happy faces, respectively). Overall, information from multiple areas of the face and multiple color channels appeared to contribute to reconstruction success. For instance, the lightness of the cheeks along with the color of the forehead, especially as encoded in the red-green channel, appeared to be correctly recovered. As expected, and in agreement with the correlation results above, fMRI heatmaps evinced lower levels of accuracy while TO heatmaps evinced the highest overall accuracy.

**Figure 5 with 1 supplement.**
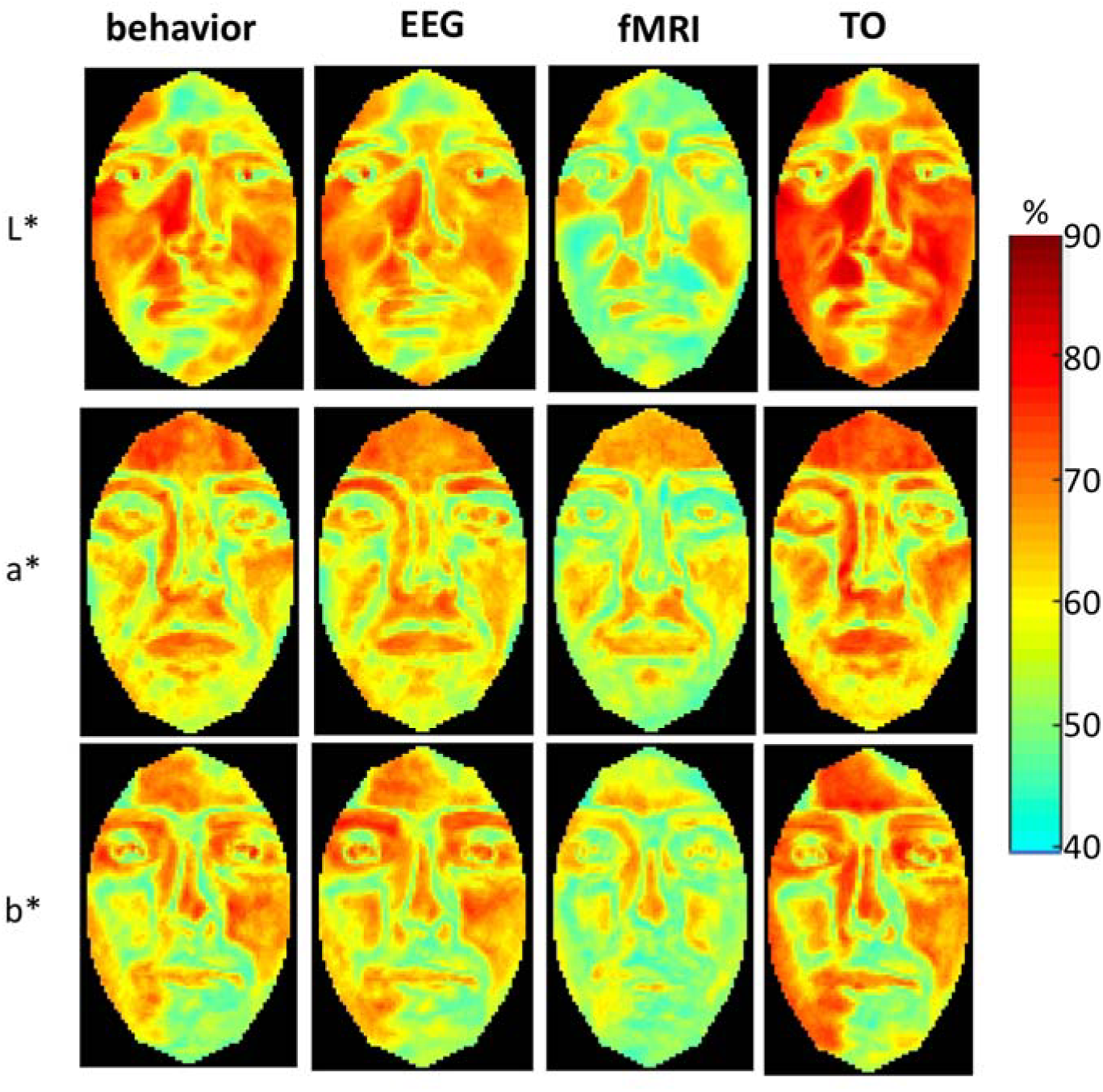
Accuracy heatmaps of surface reconstruction for neutral faces across pixels and color channels. Color indicates average reconstruction accuracy across 54 facial identities. Multiple areas of the face and multiple color channels provide accurate information for reconstruction purposes; differences in both global and local accuracy can be noticed across modalities. Figure 5 – Figure supplement 1. Accuracy heatmaps of surface reconstruction for happy faces across pixels and color channels.

### Facial image reconstruction

Shape and surface information, as retrieved separately from each data type, was combined into recomposed image reconstructions – for examples see Figure 6. Reconstructions appeared to capture, for any given modality, visual properties indicative of facial identity.

**Figure 6.**
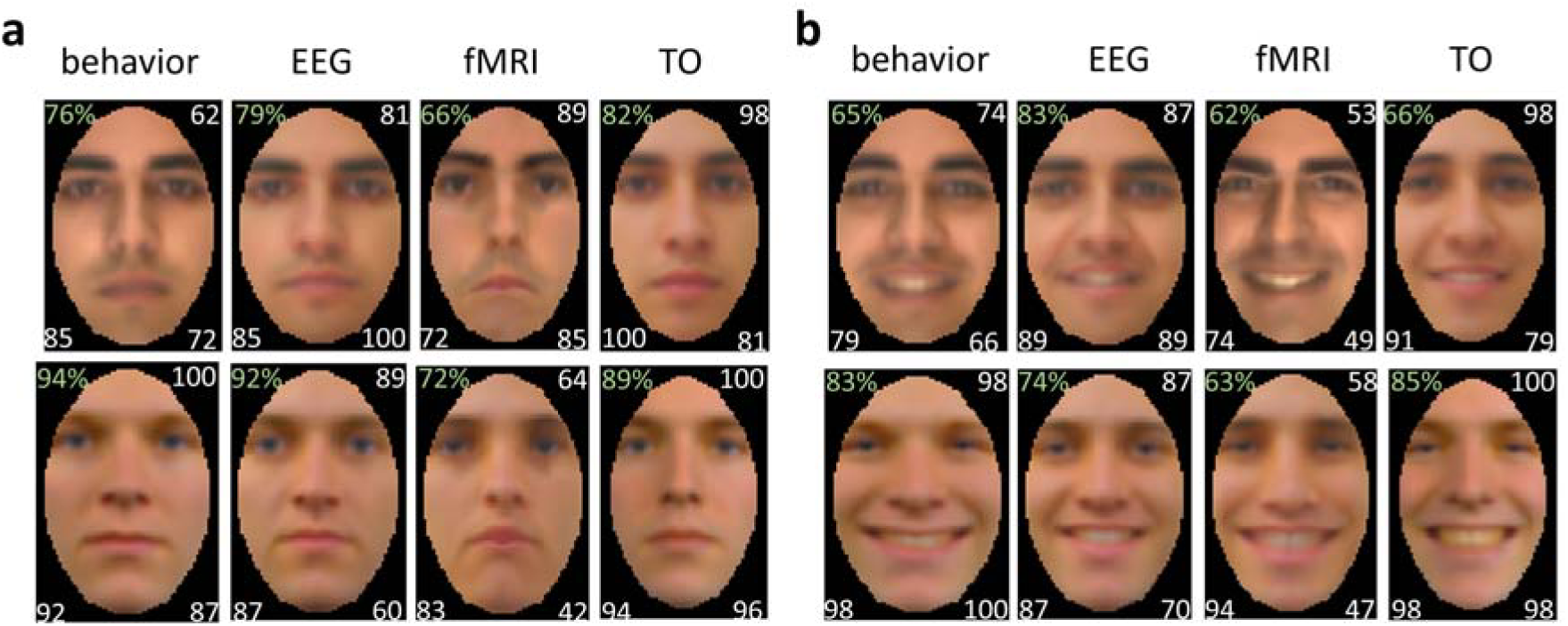
Examples of face image reconstructions (numbers in the upper left indicate experimental-based estimates of reconstruction accuracy; other image-based accuracy estimates are displayed for recomposed faces in the top right, for shape in the bottom left and for surface in the bottom right corners).* *Images of face stimuli could not be reproduced due to copyright restrictions

To evaluate this claim, empirical data were collected from a novel group of naïve observers who matched reconstructions against corresponding stimuli. Performance (Figure 7) was well above chance for all modalities and expressions (*p*’s<0.001; one-sample two-tailed t-test against 50% chance performance in a 2AFC task; Bonferroni correction across comparisons). A two-way repeated measures analysis (4 modalities x 2 expressions) found, as expected, a main effect of modality (F(3,75)=18.30, *p*<0.001, *η^2^*=0.196) but no effect of expression (*p*=0.140) and no interaction (*p*=0.333). While a direct comparison of empirical modalities in terms of accuracy may be misleading, as discussed above, it is of interest to assess how closely empirical modalities can approach the level of TO performance. Planned comparisons between TO and each empirical modality, collapsed across expressions, showed that TO surpassed fMRI (*p*<0.001), but not EEG (*p*>0.556) or behavioral data, which provided marginally better results (*p*=0.067).

**Figure 7.**
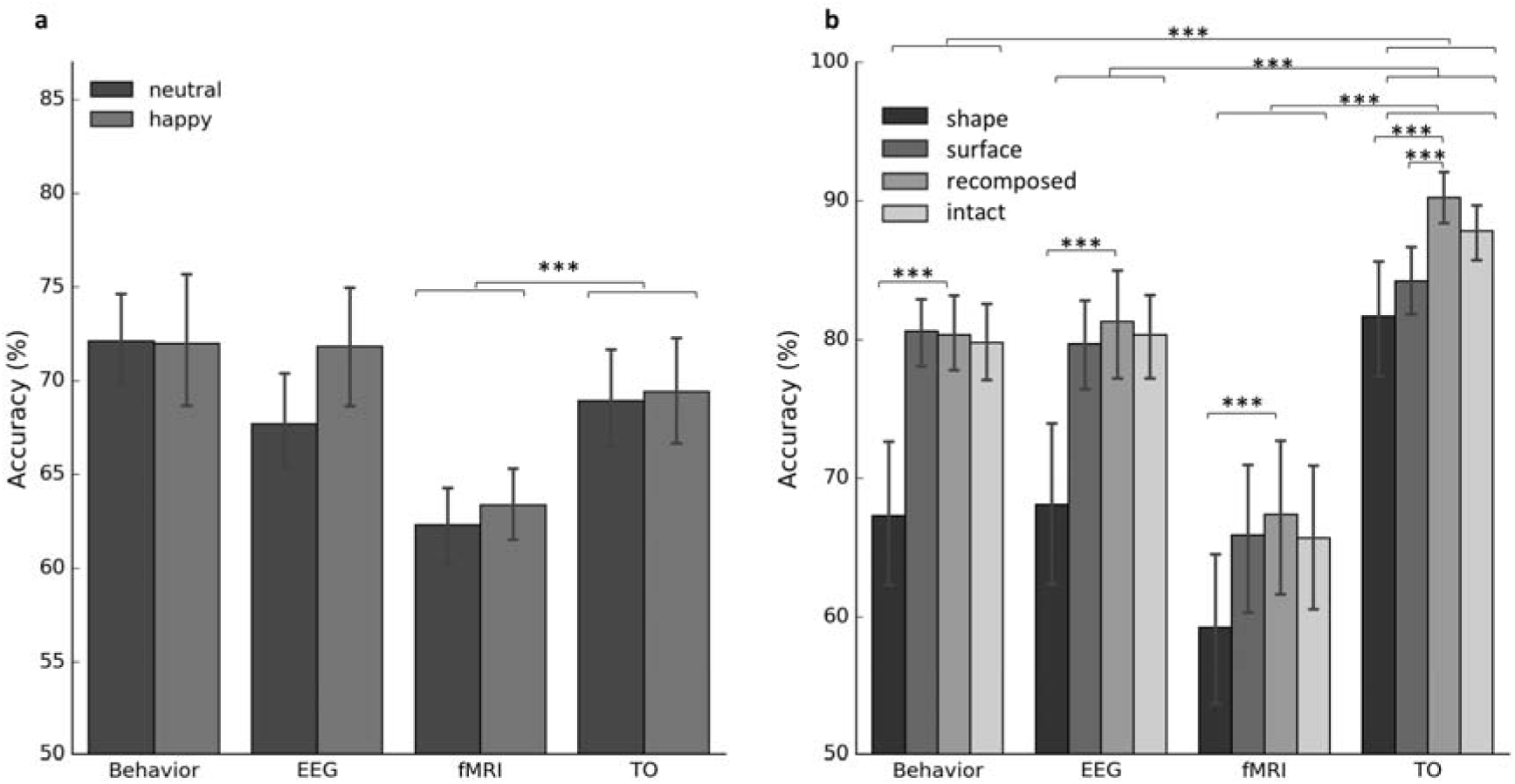
Reconstruction accuracy based on (a) experimental estimates and (b) image-based estimates. The outcome of planned comparisons is shown (a) between TO and other modalities as well as (b) between recomposed faces and other types of reconstruction (*** *p*<0.001). Error bars indicate ±1SE (a) across participants and (b) across items.

Next, we assessed whether the combination of shape and surface provides any advantage over reconstructed shapes and surface in isolation, as well as over a previous version of reconstruction that does not appeal to shape-surface decomposition (Nestor et al, 2016), for short here ‘intact reconstructions’. To this end, we considered objective estimates of reconstruction accuracy (Figure 7b): for recomposed faces this was computed across pixel intensities in the same manner in which surface accuracy was derived. Since permutation tests were not feasible for reconstructed faces, as they rely on the manual combination of shape and surface information (see Materials and Methods), parametric tests were conducted here across items (e.g., 54 facial identities). We note that while parametric tests are less conservative than the permutation tests above, their goal here is not to assess performance against chance but, rather, to explore differences across different types of reconstruction. A three-way analysis of variance (4 data types X 4 reconstruction types X 2 expressions) found, as expected, a main effect of data type (F(3,159)=44.00, *p*<0.001, *η^2^*=0.160), a main effect of reconstruction type (F(3,159)=25.82, *p*<0.001, *η^2^*=0.054) along with an interaction between data type and reconstruction type (F(9,477)=3.36, *p*=0.016, *η^2^*=0.008). No significant effect or interactions were found for expression (*p*>0.05). Further pairwise comparisons found that recomposed faces were reconstructed more accurately than shape in all instances (*p*’s<0.001) (Figure 7b) but only surpassed surface in the case of TO (*p*<0.001). Last, recomposed reconstructions were systematically more accurate than intact reconstructions (Nestor et al, 2016; Nemrodov et al, 2018); however, the difference did not reach significance for any modality (*p*’s>0.139).

Thus, the benefit of combining shape and surface reconstruction for reconstruction purposes is clearly apparent only for TO. This result is consistent with the less efficient retrieval of shape information from behavioral and neural data noted above and suggests comparatively higher reliance on surface information in visual face processing.

### The spatiotemporal profile of shape and surface processing

To investigate in further detail the temporal dynamics of shape and surface processing, reconstruction accuracy was computed for both types of information across occipitotemporal (OT) electrodes using a ∼10ms sliding window - the time course of reconstruction averaged across facial identities is displayed in Figure 8 and an example of recomposed reconstruction over time is shown in Movie 1. Overall, we found that surface information was more accurately retrieved than shape information, yet both evinced multiple intervals of above-chance reconstruction (two-tailed permutation test; FDR correction over time). Specifically, they both reached significance around 150ms after stimulus onset and gradually declined after 300ms.

**Figure 8.**
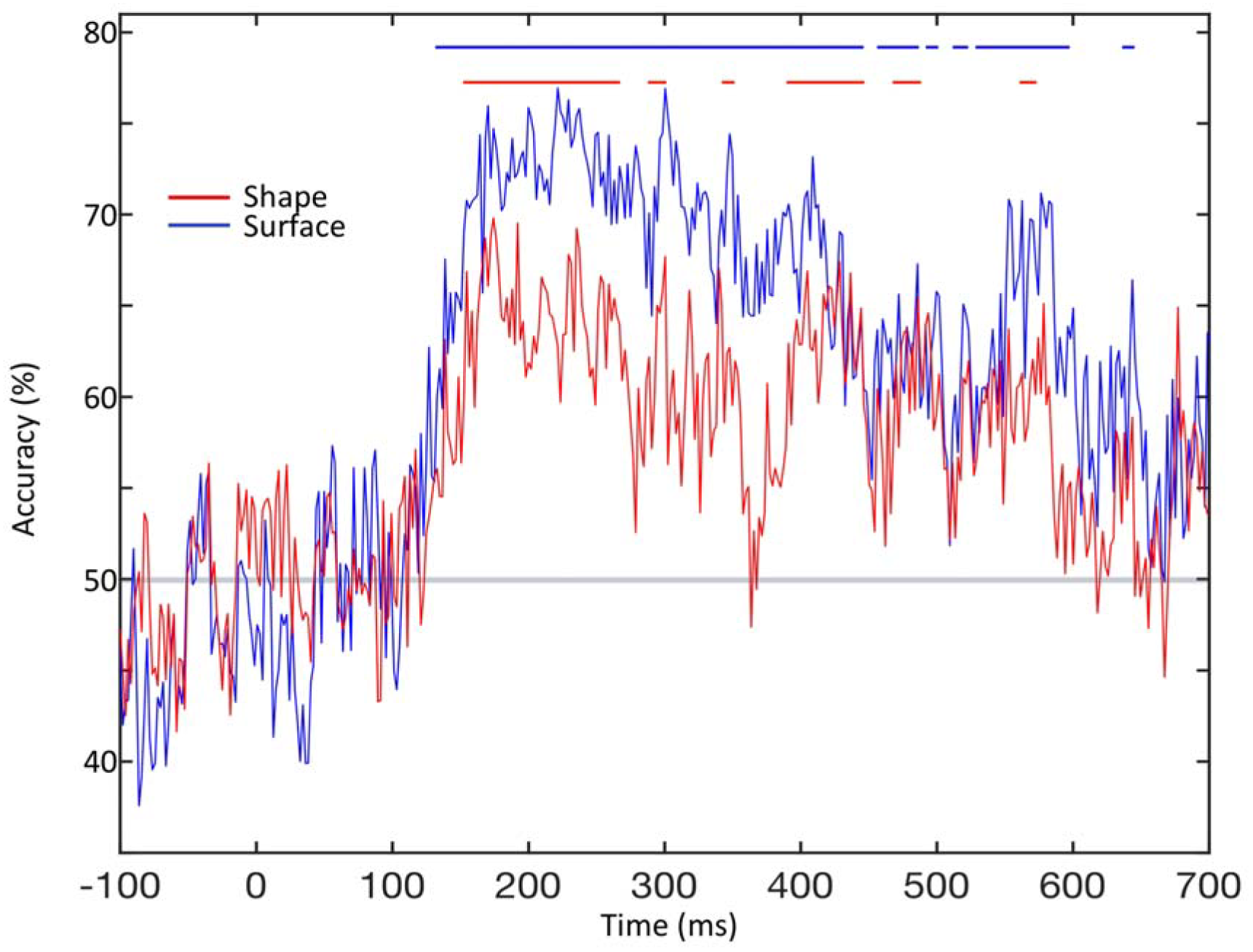
The time course of EEG-based reconstruction accuracy for shape and surface information estimated with a sliding ∼10ms temporal window. Accuracy for both types of information was above chance across multiple intervals as indicated by corresponding segments at the top of the plot (permutation test; FDR-correction across time, *q*<0.01).

To further clarify the neural locus of relevant information, reconstruction results were computed across fMRI patterns in bilateral pairs of FG areas. Specifically, to evaluate the posterior-to-anterior progression of information, reconstruction results were recomputed separately for bilateral posterior FG areas, for anterior FG areas as well as for inferior frontal gyrus areas capable of supporting face decoding (Nestor et al, 2016). Reconstruction results (Figure 9) pointed to above-chance accuracy for both shape and surface information in posterior as well as in anterior FG areas, with equivalent levels of accuracy across regions, but not in IFG areas (two-tailed permutation test; Bonferroni correction).

**Figure 9.**
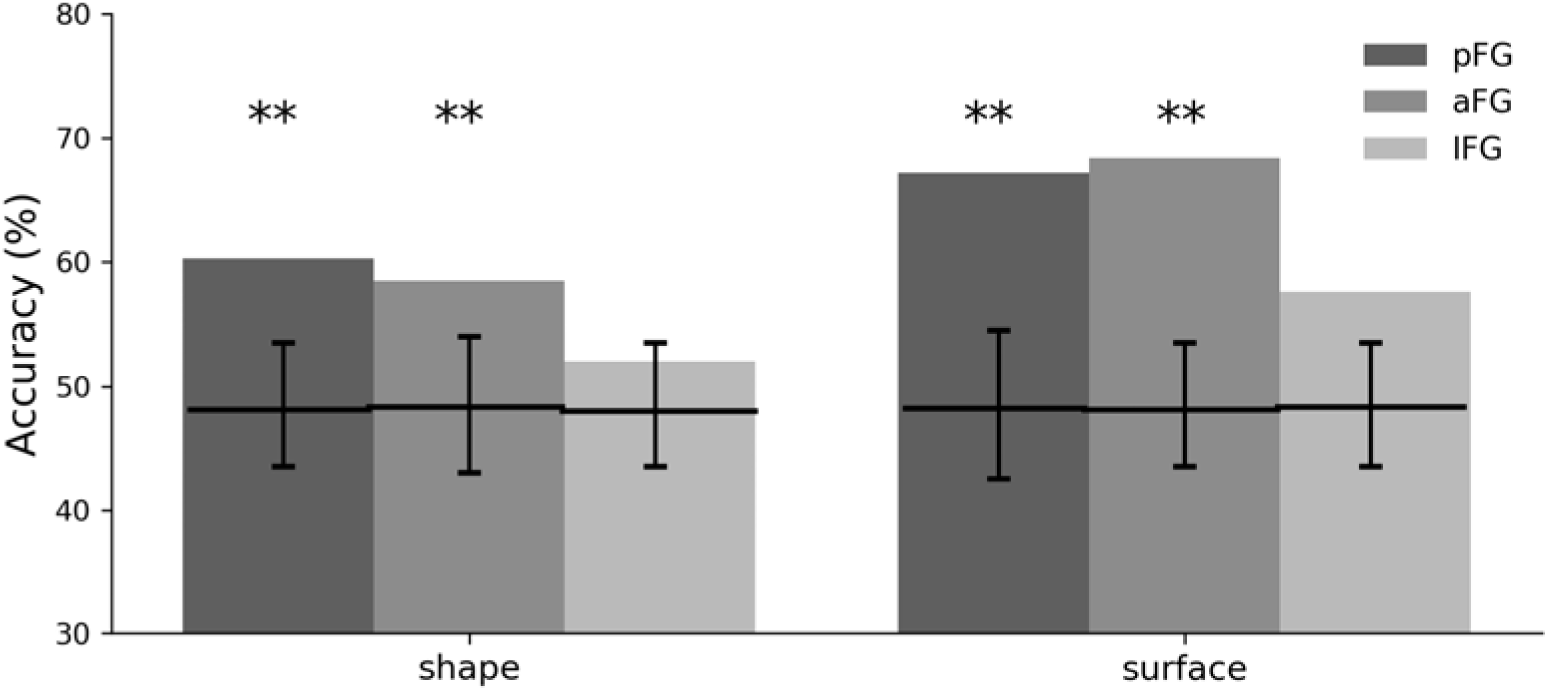
Reconstruction accuracy for three bilateral ROIs. Fusiform gyrus areas but not inferior frontal gyrus areas supported above-chance reconstruction of shape and surface (*** *p*<0.001, two-tailed permutation tests; Bonferroni correction).

Last, to relate the temporal and the spatial profile of reconstruction-relevant information, accuracy estimates from EEG and fMRI were correlated across time intervals and regions separately for shape and surface properties (Pearson correlation; FDR-correction across time points). In the case of shape, the results found a significant correlation between posterior FG-based estimates and EEG estimates around 180 ms after stimulus onset (Figure 10). Multiple time points around 170 ms, but also for subsequent intervals evinced significant correlations with anterior FG estimates but not with IFG ones. A similar investigation of surface information revealed smaller correlation values and no significant correlation with any ROI (Figure 10 – Figure supplement 1) in agreement with the results reported above (e.g., Fig. 2c).

**Figure 10 with 1 supplement.**
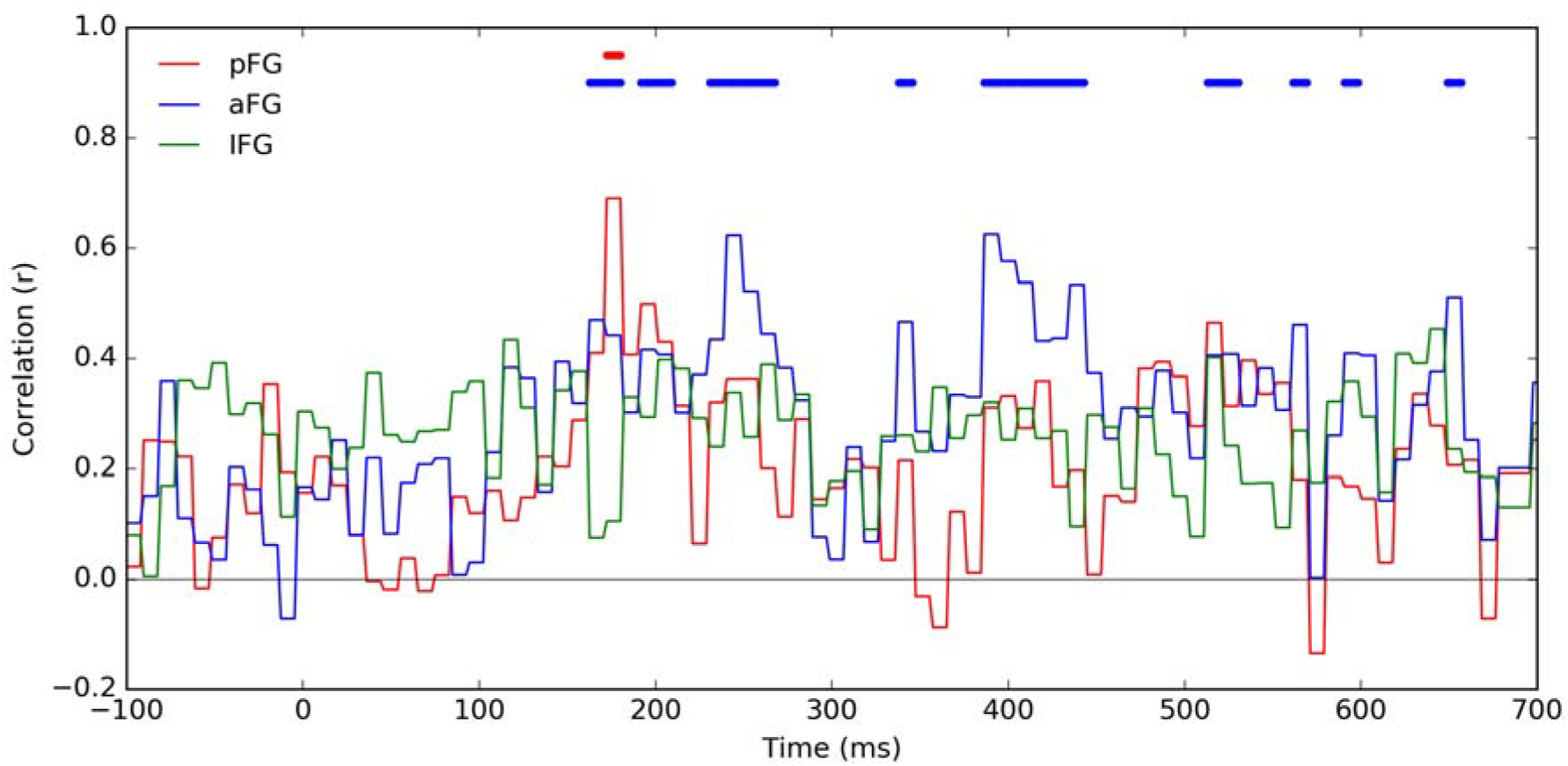
Correlation of reconstruction accuracy for shape based on fMRI data from three ROIs and from EEG data across ∼10ms temporal intervals. Results are shown for shape. Intervals of significance are marked by corresponding segments at the top of the plot (Pearson correlation; FDR correction across time, *q*<0.01). Figure 10 – supplement figure 1. Correlation of surface reconstruction accuracy for surface based on fMRI data from three ROIs and from EEG data across ∼10ms temporal intervals. No interval reaches significance (Pearson correlation; FDR correction across time, *q*<0.01).

## Discussion

The present study examined the representational basis of shape and surface information underlying individual face processing. This investigation capitalized on a robust approach to image reconstruction to uncover and relate relevant representational structures captured by distinct modalities. The ability to retrieve such information successfully and consistently enabled us to address a number of key questions as follows.

First, we examined the possibility of reconstructing both shape and surface information from each of four data types: behavioral, EEG, fMRI and TO. Our results confirmed that this is indeed possible while also revealing the advantage of surface over shape for face representations. Specifically, surface information was reconstructed more accurately than shape information for each of the empirical modalities but not for an image-based TO that yielded equivalent estimates of reconstruction accuracy for the two. Also, surface information was recovered with equivalent levels of success across different color channels, speaking to the role of chromatic information in face representations. The relative contribution of shape and surface properties continues to be contested. For example, the dominant role of surface properties in face recognition has been extensively documented for familiar faces (Burton et al., 2005; Calder et al., 2001; Hancock, Burton, & Bruce, 1996; Kaufmann & Schweinberger, 2008; Russell et al., 2006; Vuong et al., 2005) while such a role has been assumed by shape properties in the case of unfamiliar faces (Jiang, Blanz, & Rossion, 2011; Lai, Oruc, & Barton, 2013). The importance of shape is also consistent with the value of configural information for holistic face perception (Leder, & Carbon, 2006; Maurer, Le Grand, & Mondloch, 2002; McKone & Yovel, 2009; Piepers & Robbins, 2012; Richler et al., 2009; Tanaka & Gordon, 2011). Yet, other evidence suggests that even for unfamiliar faces, surface could provide dominant cues (Itz, Golle et al., 2017; Russell, Sinha, Biederman, & Nederhouser, 2006) and that configural shape information is of limited use (Taschereau-Dumouchel, Rossion, Schyns, & Gosselin, 2010). In agreement with such evidence, we find that shape information is relatively underrepresented compared to its surface counterpart for unfamiliar faces. As a caveat to this conclusion, we note that the present investigation did not consider three-dimensional face shape (Jiang, Blanz & O’Toole, 2009) but only two-dimensional information – additional 3D cues may facilitate the recovery of overall shape information and lead to more precise representations (at least as accurately as that of surface properties). Such an outcome would be especially relevant for face perception in more naturalistic settings.

Second, we aimed to characterize the spatiotemporal profile of shape and surface processing. With regard to temporal dynamics, previous work targeting specific ERP components has yielded rather inconsistent results. For instance, sensitivity to the shape of unfamiliar faces has been found at the latency of the N170 ERP component (Caharel, Jiang, Blanz, & Rossion, 2009) but also earlier, for P1 (Itz, Schweinberger, & Kaufmann, 2016), and, in the case of familiar faces, later for P200 (Itz, Schweinberger, Schulz, & Kaufmann, 2014). Similarly, surface processing is apparent first at the latency of the N250 component for both familiar and unfamiliar faces (Caharel et al., 2009; Itz, Schweinberger, Schulz, & Kaufmann, 2014), yet other studies have also found N170 sensitivity to facial surface (Balas & Nelson, 2010; Brebner et al, 2011; Minami, Nakajima, Changvisommid, & Nakauch, 2015). Unlike previous work, the present investigation used pattern analysis applied to entire epochs rather than univariate analyses targeting specific ERP components. Following this approach, we found that reconstruction accuracy reaches significance for both shape and surface around 150ms and exhibits an extended interval of above-chance performance. From a theoretical standpoint, we note that a similar time course for shape and surface processing could prove advantageous for the efficient integration of this information into unified face percepts. Also, the present results are in broad agreement with the presence of facial identity information at the latency of N170 (Caharel et al., 2009; Itier & Taylor, 2002; Nemrodov et al., 2016).

Regarding the cortical locus of shape and surface processing, previous work has found sensitivity to shape (Gao & Wilson, 2013; Gilaie-Dotan, Gelbard-Sagiv, & Malach, 2010) and surface (Harris, Young, & Andrews, 2014; Jiang et al., 2009) information within the fusiform face area (FFA), consistent with the notion that this region contains unified representations of faces (Liu, Harris, & Kanwisher, 2010). Also, recent research has found equivalent adaptation effects to shape and surface information in the occipital face area (OFA) and in the FFA (Andrews et al., 2016). While our investigation targeted regions localized through pattern analysis rather than face-selective regions per se, our results also support the idea of shape and surface integration within common regions subserving facial identity representations. Specifically, shape and surface information was recovered from posterior and anterior fusiform areas able to discriminate different facial identities. In contrast, another IFG region capable of such discrimination was unable to support either shape or surface reconstruction. Interestingly, a frontal area involved in face processing has been found in the human and monkey brain (Axelrod & Yovel, 2015; Rajimehr, Young & Tootell, 2009; Tsao et al., 2008). Recent work has argued that this area hosts higher-level, view-invariant facial representations (Guntupalli, Wheeler & Gobbini, 2017) facilitating access to person knowledge through the extended system for face perception (Collins & Olson, 2014; Haxby et al, 2000). This possibility accounts for the inability of the IFG to support image reconstruction, as reported above, while also pointing to the need to characterize more precisely the transformation of visual information across a hierarchy of face processing regions.

Third, we assessed the correspondence of facial information retrieved by different modalities. A widely influential approach, representational similarity analysis (Kriegeskorte, Mur & Bandettini, 2008), has been instrumental in relating representational structures captured by different neuroimaging modalities and computational models (Carlin & Kriegeskorte, 2017; Carlson et al., 2013; Cichy, Pantazis, & Oliva, 2016; Olander et al., 2017). Overall, this approach has uncovered both commonalities and complementarity in the visual information retrieved by different modalities (Cichy, Pantazis & Oliva, 2016). Our results agree with this conclusion while further analyzing the source of common/distinct visual information in terms of shape and surface. Specifically, RSA revealed overall similarity of representational structure for faces across different modalities. However, a more detailed investigation of reconstructed information showed that shape is more consistently recovered across modalities relative to surface dimensions. For instance, fMRI yielded surface reconstructions that did not match well their behavioral and EEG counterparts. Yet, this finding could be due to the fact that the fMRI signal considered reflects different processing stages of facial information relative to other modalities.

To address this possibility, reconstruction accuracy was systematically related across different intervals and areas. Interestingly, in the case of shape, this analysis revealed significant correlations between EEG-based results and their fMRI counterpart. Specifically, such correlations were noted as early as 170ms after stimulus onset for both posterior and anterior FG areas. However, these correlations only persisted at later latencies for aFG areas consistent with a spatiotemporal hierarchy of processing steps in face perception. In contrast, in the case of surface, EEG and fMRI yielded lower, non-significant correlations suggesting partly different representations recovered by these two neuroimaging modalities.

At a finer-grained level, the issues discussed above emphasize the need to elucidate the specific features underlying face processing and their neural representations. An evaluation of reconstruction heatmaps show that eye shape and color information is especially well retrieved in agreement with the role of such information for face recognition (Ince et al., 2016; Issa & DiCarlo, 2012; Nestor, Vettel & Tarr, 2008). Additional information regarding nose and mouth shape could also be retrieved (Abudarham & Yovel, 2016) while forehead and cheek colors were recovered with various degrees of accuracy across different color channels. Importantly, such cues appeared to reflect objective information content as revealed by TO reconstructions. Further, heatmaps of lighting and color channels exhibited different spatial patterns suggesting that chromatic information supplements lighting-based representations of facial identity (Nestor et al, 2013). Chromatic information stored in high-level visual areas may facilitate such representations and account for the proximity of these areas to face-processing cortex (Lafer-Souza et al, 2016).

While the present investigation targets the representational basis of shape and surface processing via image reconstruction, we also note here the converse aspect of this investigation. Specifically, we inquire into the benefit of shape and surface decomposition for image reconstruction purposes. Previous work has relied on the coarse alignment of facial features to minimize the need for such decomposition (Cowen, Chun, & Kuhl, 2014; Nestor, Plaut, & Behrmann, 2016; but see Chang & Tsao, 2017; Zhan et al., 2017). Hence, it is useful to assess, in the context of the present investigation, whether recomposed faces are more accurately reconstructed than intact ones. Interestingly, decomposition appeared to provide a systematic advantage to reconstruction across modalities, yet this advantage did not reach significance in any given case. Clearly, shape-surface decomposition would be required when dealing with pronounced image variability such as that due to viewpoint. However, if such variability is controlled across stimuli and if feature alignment is successfully carried out in advance it appears that shape-surface decomposition confers, at most, a minimum advantage to reconstruction.

Of relevance here, we note that shape-surface manipulations introduce a systematic loss of information due to image (re)warping and that this loss is likely to limit reconstruction success (see Materials and Methods, Stimulus shape-surface re/re-composition). To address this and, also, to allow the extension of current work to a wider, more diverse class of face images, state-of-the-art algorithms for fiducial point detection and shape analysis could be employed (Ahdid, Taifi, Safi & Manaut, 2016). Such methods would confer robustness to shape-surface decomposition and facilitate the integration of reconstruction results with algorithms for face recognition that rely on elaborate schemas for shape/surface analysis (Zhao, Chellappa, Phillips & Rosenfeld, 2003) – such integration could serve specific goals for translational research (e.g., the automatic identification of a face image reconstructed from eyewitness memory).

At the other end, we note certain theoretical limitations associated with the use of naturalistic stimuli. Of specific relevance here is the covariation of different types of information conveyed by shape and surface properties. For instance, shading as captured by surfaces can provide cues to 3D shape (Kramer, Jenkins, & Burton, 2016). Such covariation constraints the interpretation of our results in important ways. Concretely, in the context of empirical modalities, we found no advantage to combining shape and surface information over using surface information in isolation (though such an advantage was noticed for TO). The lack of this advantage could simply reflect the covariation of shape and surface properties in natural images. Thus, to better disentangle the distinct contribution of such properties to face representations, one may be better served by artificial stimuli that manipulate orthogonally distinct types of information and, also, by image processing techniques that ascribe 3D cues exclusively to shape (O’Toole et al, 1999; Paysan, Knothe, Amberg, Romdhani, Vetter, 2009).

In summary, the present work seeks to uncover the representational underpinnings of shape and surface processing in face perception through the use of novel image-reconstruction methodology. Our results show that such information can be reliably extracted from multiple modalities, that its representational structure is partly shared across modalities and that its spatiotemporal profile speaks to the close integration of shape and surface cues in face processing. More generally, the present findings showcase the value of image reconstruction methodology in elucidating the content and the neural profile of visual representations.

## Materials and methods

### Stimuli

A common subset of 108 stimulus images was identified across three different studies investigating empirical and computational aspects of unfamiliar face recognition (see Experimental procedures). Images of 54 individuals displaying neutral and happy facial expressions were selected from three databases: AR (Martinez & Benavente, 1998), FEI (Thomaz & Giraldi, 2010) and Radboud (Langner et al., 2010). All images featured young adult Caucasian males with frontal view, gaze and illumination. The stimuli were selected so that no facial accessories, hair or makeup obscured the internal features of the face and so that all happy expressions displayed an open-mouth smile. These images were: (a) scaled uniformly and aligned with roughly the same position of the eyes and the nose; (b) cropped to eliminate background; (c) normalized with the same mean and root mean square (RMS) contrast values separately for each color channel in CIEL*a*b* color space, and (d) reduced to the same size (95 X 64 pixels). Note that this procedure did not change the aspect ratio of the images though the position of the eyes and the nose was roughly the same across stimuli. Thus, every effort was made to homogenize the stimulus set both in terms of low-level and high-level face properties preventing the potential contribution of such factors to image reconstruction.

### Participants

All participants (age range across studies: 18-34 years; 21 males, 22 females) were Caucasian adults with normal or corrected-to-normal vision and no history of cognitive or neurological disorder. All participants provided informed consent and all experimental procedures were approved by the Research Ethics Board at University of Toronto and/or the Institutional Review Board at Carnegie Mellon University.

### Experimental procedures

Data used for reconstruction purposes were selected from three previous studies as follows.

Behavioral data consisted of similarity ratings with pairs of faces acquired from 22 participants (reported in Nestor et al, 2013, Experiment 1). Briefly, on each trial, participants were presented with two facial identities, one neutral and one happy, side by side, for 400ms, and were asked to judge their visual similarity on a 5-point scale. Each participant rated all possible 1431 facial pairs, corresponding to 54 facial identities - for clarity, only a subset of the original data were considered here (i.e., 6 additional facial identities were not used in the EEG study summarized below and, hence, were excluded from further analyses of behavioral data).

EEG data were previously acquired from 13 participants who performed a go/no-go gender categorization task (Nemrodov et al, 2018). On ‘no-go’ trials, participants viewed the stimuli described above while, on ‘go’ trials, they were asked to press a designated key in response to the appearance of a female face. Each of the 108 main stimuli was presented for 300 ms and repeated across 64 trials for each participant.

fMRI data were acquired from 8 participants who performed a continuous one-back identity task (Nestor et al, 2016). Briefly, on each trial, participants viewed a stimulus for 900 ms and responded whether the current stimulus displayed the same individual as that presented on the previous trial, irrespective of emotional expression. The experiment used a wide-spaced design (8s trials) and allowed for the repetition of each stimulus for a minimum of 10 trials across five 1-hr sessions for each participant. Again, only a subset of the stimuli used in the original study is considered here to enable direct comparison with data from the other modalities.

To be clear, we note that the neuroimaging studies above (Nemrodov et al, 2018; Nestor, Plaut & Behrmann, 2016) did not separate shape and surface cues for reconstruction purposes nor did they assess the contribution of such cues to visual face representations. Further, the behavioral study above (Nestor et al, 2013) did not target any form of image reconstruction and, thus, it provides a new testing ground for reconstruction endeavors.

### Representational similarity analyses

Our reconstruction procedure fundamentally relies on the structure of representational (dis)similarity matrices (Kriegeskorte, Mur, & Bandettini, 2008) to derive facial image features and to use such features for reconstruction purposes. Hence, the first step of our investigation is to construct such matrices separately for each data type.

Specifically, for each modality and for each participant, a similarity matrix was designed to store pairwise similarity estimates across 54 facial identities. In the case of behavioral data, these estimates were readily available in the form of similarity ratings. In the case of EEG and fMRI data such estimates were derived through one-against-one pattern classification of different identities, separately for each expression, using linear support vector machines (SVM). Briefly, pairwise classification was applied across EEG spatiotemporal patterns recorded across at 12 occipitotemporal (OT) electrodes (left: P5, P7, P9, PO3, PO7, O1; right: P6, P8, P10, PO4, PO8, O2) during an interval spanning 50-650ms after stimulus onset – these spatiotemporal patterns were selected based on their ability to support face decoding (Nemrodov et al, 2018). Analogously, for fMRI, classification was applied across multivoxel patterns within areas supporting above-chance face discrimination as identified through prior searchlight mapping. Such areas were previously identified (Nestor et al, 2016) bilaterally in the posterior fusiform gyrus (group-map Talairach coordinates: left FG, –39, –49, –16; right posterior FG, 39, −69, −4), the anterior fusiform gyrus (left FG, –34, –36, –14; right anterior FG/parahippocampal gyrus, 31, −19, −9) and the inferior frontal gyrus (left IFG, −36, 24, −9; right IFG, 46, 11, −1). Thus, discrimination accuracy computed across neural patterns in these regions was used to estimate the similarity of the stimuli eliciting such patterns – a more detailed account of data preprocessing and pattern analyses can be found in the studies above.

In addition, objective measures of image similarity in CIEL*a*b* color space were computed for the purpose of constructing a theoretical observer (TO) exploiting low-level visual similarity. To this end, pixelwise Euclidean distances were computed across all pairs of facial identities separately for each expression and the results were stored in corresponding similarity matrices. Of note, while more elaborate models of face similarity are of interest (e.g., Carlin & Kriegeskorte, 2017), the TO above is particularly relevant given that pixelwise similarity is also one of the main criteria for assessing reconstruction quality. Thus, a TO matching typical criteria for result assessment is particularly well-suited for estimating an upper limit of reconstruction success.

Last, similarity estimates were averaged across participants and across expressions to deliver a single similarity matrix for each modality: behavioral, EEG, fMRI and TO. These resulting estimates were related across modalities via Spearman correlation to estimate the presence of a common representational structure.

### Stimulus shape-surface de/re-composition

All stimuli were tested for their ability to undergo reliable shape-surface decomposition and recomposition. Specifically, for reconstruction purposes, all stimuli were analyzed into shapes (i.e., configurations of fiducial points, such as the corners of the eyes and the tip of the nose, labeled with their geometric coordinates) and shape-free surfaces (i.e., facial images warped to a common shape template). To this end, fiducial points were manually marked for each stimulus using the Interface toolbox (Kramer, Jenkins, & Burton, 2016) and, then, the marked stimulus was warped to a preset shape template. Thus, the shape of each stimulus is represented as a vector of fiducial point coordinates (82 points x 2 in-plane coordinates) while its surface is represented by a template-warped image.

The process above was then reversed by recombining shapes and surfaces into approximations of the original stimuli. This procedure was carried out to estimate information loss inherent to de/re-composition due to image (re)warping and, thus, to assess, the objective cost of shape/surface manipulations for reconstruction. To this end, rewarped versions of the stimuli obtained through recomposition were compared against actual stimuli. Concretely, for each rewarped stimulus we computed the ratio between the pixelwise Euclidean distance relative to its original version and the distance to every other stimulus, one at a time; hence, ratios larger than 1 would render rewarped stimuli more similar to other facial identities. The outcome of these computations (mean ± 1SD across 54 identities) yielded ratios of 0.333 (±0.054) and 0.329 (±0.052) for neutral and happy faces, respectively. Thus, shape-surface de/re-composition generally preserves identity information but it does introduce systematic image distortions likely to limit reconstruction success.

### Reconstruction procedure

The current procedure builds upon previous work (Nestor et al, 2016; Nemrodov et al, 2018) aimed at deriving pictorial features directly from the structure of empirical data and to use them for facial image reconstruction. Here, we further develop this procedure to derive separate sets of shape and surface features from multiple data types and we assess their relative contribution to face representations as revealed by image reconstruction.

For each modality, the reconstruction procedure was separately conducted following a sequence of steps (Figure 1). Concisely, in order to reconstruct the appearance of any given target, the procedure involved: (i) selecting a similarity submatrix containing the pairwise similarity of all faces other than the target; (ii) estimating the dimensions that structure face space by applying multidimensional scaling (MDS) to the resulting submatrix; (iii) deriving, for each dimension, shape and surface features through a strategy akin to reverse correlation; (iv) assessing feature significance and selecting a subset of informative features; (v) projecting the target face into the existing face space based on its similarity with the other faces; (vi) reconstructing the shape and the surface of the target face through a linear combination of informative features, and (vii) combining the resulting shape and surface into a single image reconstruction of the target face.

In more detail, the leave-one-out procedure enforces non-circularity by excluding the reconstruction target from the estimation of face space and its underlying features. Specifically, a face space construct was derived from the pairwise similarity of 53 facial identities and, then, its corresponding features were used in the reconstruction of the target face. To this end, a 20-dimensional face space was estimated through metric MDS, given that this number of dimensions accounted for more than 90% of data variance for any modality and, also, that it agrees with previous estimates of face space dimensionality in human recognition (i.e., 15-22) (Lewis, 2004). Then, a corresponding number of shape and surface features were computed for each dimension through an approach akin to reverse correlation/image classification (see (Murray, 2011) for a review). Notably, this approach aims to synthesize facial features responsible for face space topography through a linear combination of face properties (i.e., shape vectors or surface images). This combination was computed as a sum of shapes and surfaces for all faces weighted by the z-scored coordinates of the corresponding faces on any given dimension. Thus, the outcome of these computations delivers, for each dimension, one shape feature, or ‘classification vector’ (CV), and one surface feature, or ‘classification image’ (CIM) – for clarity, each surface feature consists in a triplet of images, one for each color channel in CIEL*a*b*.

Further, we considered the possibility that not all face space dimensions encode visual information (e.g., as opposed to higher-level semantic information or just noise). Also, it is possible that sources of shape and surface information are differently distributed across dimensions and their corresponding features. Hence, it is important to identify relevant subsets of features that can contribute meaningful information to reconstruction. To this end, each feature was assessed for the presence of significant information. Specifically, all face space identities were randomly shuffled with respect to their coordinates on each dimension and a corresponding feature was recomputed for a total of 10^3^ permutations. Then, each true feature was compared to all permutation-based features, fiducial point by fiducial point, in the case of shape, or pixel by pixel, for each CIEL*a*b* color channel, in the case of surface (two-tailed permutation test; FDR correction across points and pixels, respectively; *q* < 0.1). Following this procedure, only features that contained significant shape or surface information were selected for reconstruction purposes.

Next, the target face was projected into the existing face space. To this end, a new MDS solution was constructed for all 54 identities and aligned with the original one via Procrustes analysis using the 53 common identities between the two spaces. The resulting alignment provides us with a mapping between the two spaces that allows us to project the target face and to retrieve its coordinates in the original space. Then, informative features were linearly combined proportionally to the coordinates of the target face on each corresponding dimension and their sum was added to the average shape and surface of the 53 faces used for feature derivation. We note that face space was uniformly scaled under the constraint that all reconstructed surfaces should have the same RMS contrast and mean value in each color channel as the surfaces of the experimental stimuli – these values were equated across experimental stimuli in an effort to minimize the contribution of low-level images differences to perception (see Stimuli above). A similar manipulation was also conducted for shape reconstructions to ensure that the variance of fiducial point coordinates matched that of the stimulus shapes. Last, the shape and surface thus computed were manually combined using the Interface toolbox into a single reconstruction to which we refer as a ‘recomposed face’.

For clarity, face space is constructed here across facial identities irrespective of emotional expression (e.g., by averaging similarity matrices for the two expressions). However, reconstruction proceeds by deriving and combining features separately for neutral and happy faces (i.e., a neutral face is reconstructed from features derived from other neutral faces while a happy face is reconstructed from features derived from other happy faces). Another possibility would be to consider separate spaces for each expression; however, previous investigations evaluated and confirmed the invariance of neural-based face space across these two emotions (Nemrodov et al, 2018).

Finally, to evaluate the benefit of considering separately shape and surface information for reconstruction purposes, another set of image reconstructions were computed without appeal to this decomposition. Specifically, face stimuli were treated in the same manner as shape-free surfaces above. We refer to the outcome of this procedure as ‘intact reconstructions’.

### Estimation of reconstruction accuracy

The accuracy of reconstruction results was assessed in two different ways: by objective image-based measures and experimentally, by additional behavioral testing.

In detail, each reconstructed shape was compared to its target via the Euclidean distance computed across corresponding fiducial point coordinates. Then, the accuracy of its reconstruction was measured as the proportion of instances for which that distance was smaller than the distance between the reconstruction and any stimulus shape other than the target. This procedure was conducted for all reconstructions, separately for each expression and each modality.

Reconstructed surface accuracy was measured similarly, except that Euclidean distances were computed across pixel values in CIEL*a*b* space. The same procedure was also followed for recomposed and intact reconstructions.

Statistical significance was then assessed through permutation tests. Specifically, the shape and the surface of each target was recomputed based on the random shuffling of identity labels across all points in face space (for a total of 10^4^ permutations). Then, the accuracy of the true reconstructions was related to that of permutation-based reconstructions (two-tailed permutation tests).

However, this procedure was not feasible in the case of recomposed faces (since that would require the manual recombination of shape and surface for a prohibitively high number of permutation-based reconstructions). For this reason, and, also, to provide a complementary evaluation of reconstruction results, additional behavioral testing was conducted as follows.

Reconstructed images consisting of 432 recomposed faces (54 identities X 2 expressions X 4 modalities: behavioral, EEG, fMRI, and TO) were judged in terms of their relative similarity to the actual stimuli. To this end, 27 new participants (seven males and twenty females, age range: 18-25) were asked to match image reconstructions to their targets in a two-alternative forced choice (2AFC) task. Specifically, each reconstruction was displayed in the presence of two stimuli, one of which was the actual target and the other another face image. Thus, on each trial, a display was shown containing a reconstructed image, at the top, and two stimuli side by side, at the bottom, all of which had the same expression and the same size (see Stimuli). Each display was presented until participants indicated which stimulus was more similar to the top image by pressing a designated left/right key. For each participant, any reconstructed image was presented 4 times along with different foils so that, across participants, each reconstruction was presented together with every possible foil. Stimulus order was pseudorandomized so that no reconstruction appeared twice on consecutive trials and target stimuli appeared equally often on the left/right side. Each participant completed 1728 trials divided equally across 9 blocks. Experimental testing was conducted within a single 1.5-hr session.

Parametric statistical analyses were next conducted to assess reconstruction success against chance (one-sample two-tailed t-tests against 50% chance-level participant performance) as well as the relative success of reconstruction across modalities and expressions (2-way factorial analysis of variance: 4 modalities X 2 expressions).

To further compare and account for the outcome of different types of reconstruction, additional analyses were conducted as detailed below.

### Evaluation of reconstruction results

First, parametric tests across items (i.e., across facial identities) were carried out to estimate the relative success of reconstruction results. Specifically, we conducted a three-way factorial analysis of variance (4 modalities X 2 expressions X 4 reconstruction types: shape, surface, recomposed and intact reconstructions) along with planned comparisons – accuracies were collapsed across expressions given the lack of evidence for a corresponding effect from prior analyses. Of note, parametric analyses across items provide a liberal way to estimate reconstruction success (e.g., compared to the permutation tests above); however, the goal of the current analysis was not to estimate significance against a preset chance level but rather to evaluate differences in reconstruction success across modalities and reconstruction types.

We note that comparing modalities in terms of overall reconstruction accuracy can be informative, by answering, for instance, how closely empirical modalities can approach TO-level performance. However, such an analysis provides an incomplete and potentially misleading picture of the relationship across empirical modalities since any difference, or lack thereof, can be the outcome of differences in experimental designs separately optimized for each modality (e.g., stimulus duration, number of stimulus presentations, task, number of participants).

Second, and aiming to address the concern above, reconstruction results were related with each other via correlation across facial identities for each pair of modalities. Specifically, image-based accuracies were related to each other across modalities, separately for shape and surface, via Pearson correlation. We note that this investigation parallels representational similarity analysis with the difference that, here, we correlate item-specific reconstruction accuracies as opposed to pairwise item similarity estimates. In addition, to clarify the relationship between behavioral and neural-based results, linear regression was used to account for behavioral reconstruction accuracies in terms of their EEG and fMRI counterparts separately for shape and surface reconstructions.

Third, to clarify the spatiotemporal profile of the information supporting shape and surface reconstruction, additional analyses were conducted across time, for EEG, and across different ROIs, for fMRI. Specifically, for the former, reconstruction was conducted across smaller 10ms temporal intervals between −100 and 700ms (i.e., across 60-dimensional vectors; 12 OT electrodes x 5 consecutive time points) providing us with the time course of reconstruction accuracy. In the case of fMRI, reconstruction was conducted for distinct pairs of bilateral ROIs in the posterior FG, the anterior FG and the IFG. Image-based accuracy was then estimated for each temporal interval and for each ROI pair and significance was estimated by two-tailed permutation tests.

## Acknowledgments.

This research was supported by the Natural Sciences and Engineering Research Council of Canada (A.N. and M.N.), by the National Institute of Health (to M.B.), and by a Research Competitiveness Fund Award (A.N.).

## Competing interests

The authors declare no competing financial or non-financial interests.

## Supplementary figures

**Figure 3 – Figure supplement 1.**
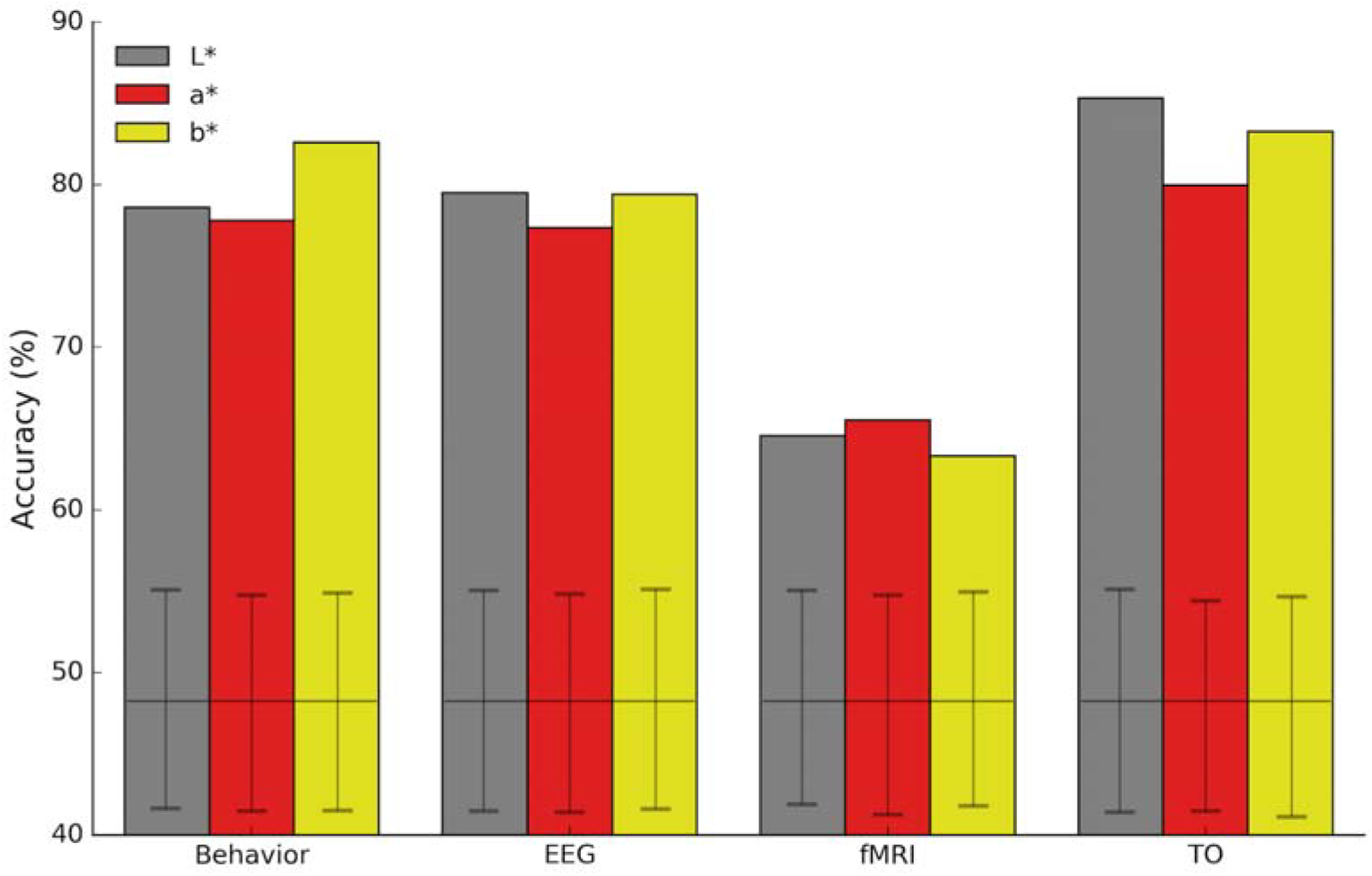
Accuracy estimates of surface reconstruction for each color channel. Results are shown for each modality collapsed across emotional expression. All estimates are above chance (all p’s<0.01, Bonferroni-corrected; two-tailed permutation tests; confidence intervals based on 10^4^ shuffles of identity labels in the corresponding face space). No difference was noticed across color channels for any modality (all p’s>0.05; two-tailed paired-comparison permutation tests).

**Figure 5 – Figure supplement 1.**
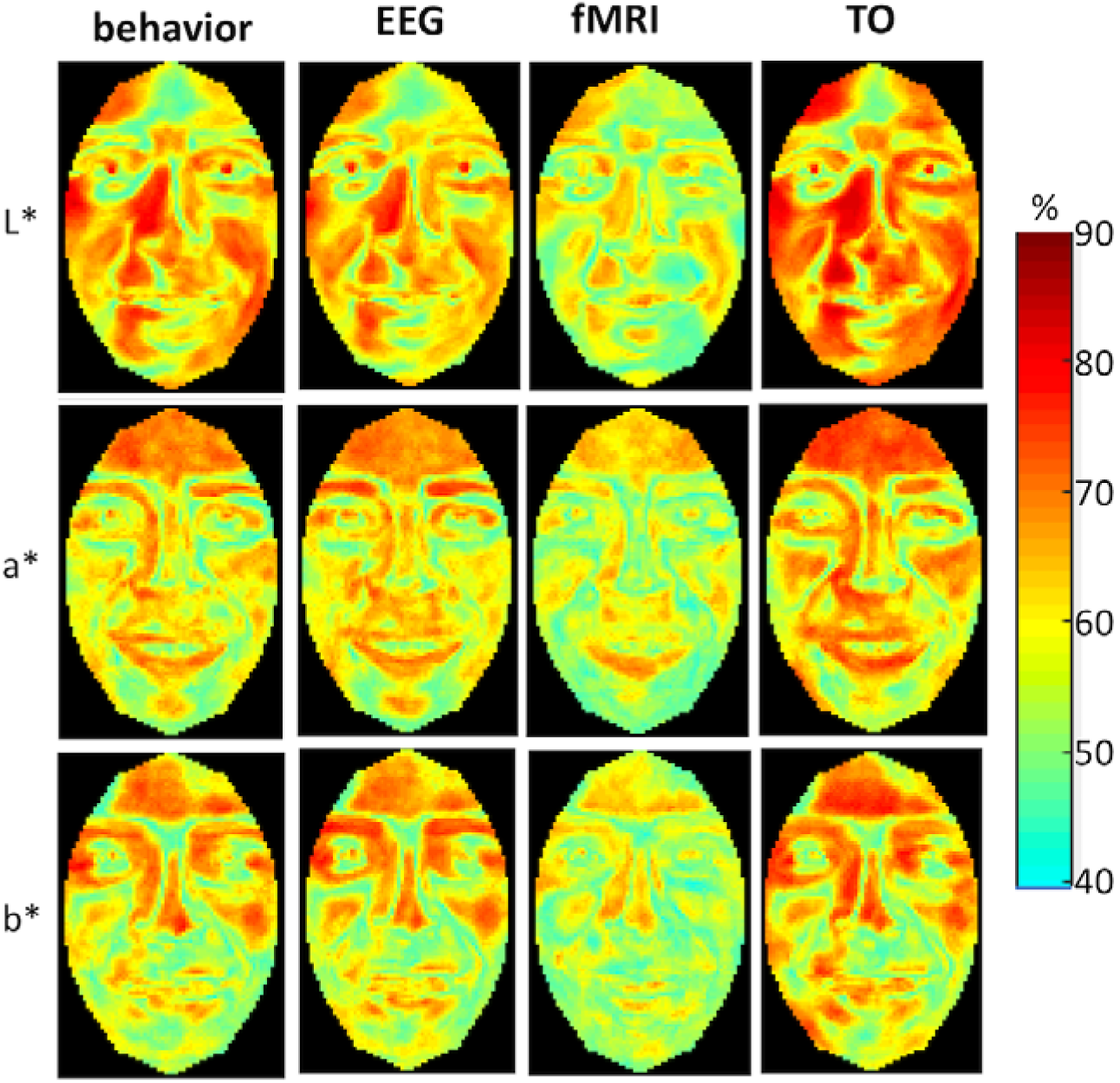
Accuracy heatmaps of surface reconstruction for happy faces across pixels and color channels. Color indicates average reconstruction accuracy across 54 facial identities. Multiple areas of the face and multiple color channels provide accurate information for reconstruction purposes; differences in both global and local accuracy can be noticed across modalities.

**Figure 10 – supplement figure 1.**
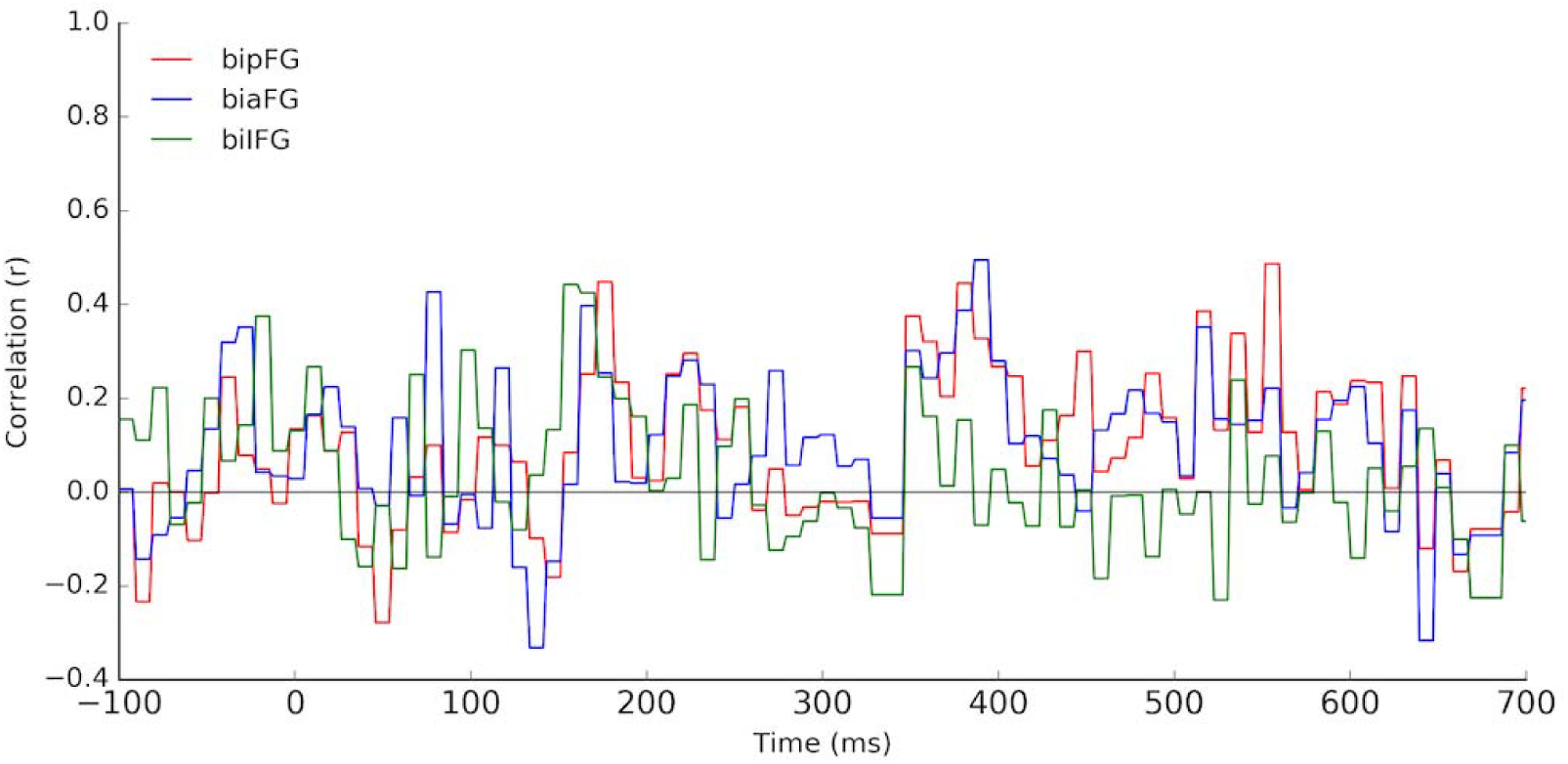
Correlation of reconstruction accuracy for surface based on fMRI data from three ROIs and from EEG data across ∼10ms temporal intervals. No interval reaches significance (Pearson correlation; FDR correction across time, *q*<0.01).

